# Systematic effects of mRNA secondary structure on gene expression and molecular function in budding yeast

**DOI:** 10.1101/138792

**Authors:** Xia Wang, Pidong Li, Ryan N. Gutenkunst

## Abstract

Dynamic control of gene expression is crucial for cellular adaptation to environmental challenges. mRNA secondary structure is known to be associated with mRNA and protein abundance, but little is known about how mRNA secondary structure affects gene expression dynamics. We report a genome-wide computational analysis of mRNA secondary structure, codon usage, and gene expression in budding yeast. We show that mRNA secondary structure combined with codon optimality regulates gene expression in multiple ways, from transcription to mRNA stability to translation. Moreover, we find that the effect of mRNA secondary structure on mRNA abundance is primarily mediated by transcription, not mRNA stability. Notably, genes with low mRNA secondary structure were substantially enriched for functions relevant to stress response, acting in the mitochondrion, endoplasmic reticulum, and ribosome. On the other hand, genes with high mRNA secondary structure were enriched for functions relevant to cellular maintenance, including macromolecular metabolism and biosynthesis. Our results suggest that mRNA secondary structure affects gene expression through coordination of multiple stages in protein biogenesis, with important consequences for stress response. The coupling of transcription to mRNA stability to translation makes concerted changes in mRNA and protein abundance possible and may amplify the effect of regulation to make quick responses to environmental variations.

## Background

Gene expression is finely and dynamically regulated through control of each step in the Central Dogma [1]. For example, recent genome-wide studies have revealed that transcription and mRNA decay rates are coupled to regulate gene expression [2-4]. In steady state, the balance between mRNA transcription and degradation determines mRNA abundance, and translation and protein degradation then impact protein abundance [5, 6]. Gene expression must also be dynamically regulated to adapt to environmental challenges, and transcription rate, translation efficiency, and decay rate limit the speed at which gene expression can change [7]. Abundant evidence shows that multiple steps in gene expression are often coordinated to shape the dynamics of the transcriptome and proteome in response to environmental perturbations [2, 4, 6, 8]. The mechanism of this coordination is, however, elusive.

Increasing evidence suggests that folding into precise secondary structures is necessary for mRNA maturation. For example, RNA secondary structures have been proposed to affect splice-site recognition [9-13] and poly(A) site recognition [12, 14]. Specific structural elements also affect the stability of many eukaryotic mRNAs [15]. In addition, RNA secondary structure elements [16, 17], such as bulge loops, hairpin loops, and stems, mediate protein-RNA interactions that play integral roles in various post-transcriptional regulatory processes, including the degradation and translation of RNA [18, 19]. RNA secondary structure thus influences nearly every step in gene expression [20-22]. Sequence features, such as codon usage, have also been proposed to affect transcription [23-25], mRNA decay [24, 26] and translation efficiency [24, 27, 28]. RNA structure is affected by codon usage [29], so RNA structure may mediate the effects of codon usage on gene expression. Recent work has shown that steady-state RNA and protein abundance are strongly correlated with RNA structure and sequence features [30, 31], and other work has delineated the effects of mRNA thermodynamic stability, sequence features, and tRNA availability on translation [32]. Nevertheless, the systematic effects of RNA structure on gene expression remain unexplored.

Recently, a high-throughput experimental approach has been developed to assay *in vitro* mRNA secondary structure. In Parallel Analysis of RNA Structure (PARS), the poly-adenylated transcript pool is treated with RNase V1 that preferentially cleaves 3′ phosphodiester bonds in double-stranded RNA and, separately, with RNase S1 that preferentially cleaves 3′ bonds in single-stranded RNA. The obtained fragments are then deep sequenced to measure the degree to which each nucleotide was in a single- or double-stranded conformation. The PARS score is defined as the log_2_ of the ratio between the number of times the nucleotide immediately downstream of a given nucleotide was observed as the first base when treated with RNase V1 and when treated with RNase S1 [33]. Techniques exist to measure *in vivo* mRNA secondary structure [12, 17, 34], which depends both on the sequence of the mRNA and on any bound proteins or RNAs. We used PARS to measure *in vitro* mRNA secondary structure to focus on the intrinsic folding of the mRNA, independent of the local cellular environment.

Large-scale manipulative experiments on RNA structure and sequence features are challenging, but the statistical technique of path analysis provides a means to disentangle the effects of multiple potential causal factors on multiple steps in gene expression from observational data [35]. Path analysis was developed to analyze complex systems in which it is challenging to isolate causes and effects, because each component potentially affects many others through a network of direct and indirect interactions. In path analysis the total correlation between any two variables is decomposed into direct effects of one on the other, indirect effects mediated by other variables, and spurious correlations due to common causes. The estimated path coefficients represent the amount of change expected in the dependent variable as a result of a unit change in the independent variable. Path analysis has been used to infer causal relationships from observational molecular data in a number of research settings, including studies of human disease [36], plant physiology [37], and protein evolution [38].

In this work, we used path analysis to decompose the correlations between mRNA structure and sequence features on transcriptional and translational processes. We then explored the functional consequences of the associations between mRNA structure and different steps of gene expression, identifying cellular functions that are enriched in genes with low or high mRNA secondary structure.

We found that mRNA structure and optimal codon usage are significantly and positively associated with transcription rate, mRNA half-life, and translation efficiency. Surprisingly, we found that mRNA secondary structure affects transcript abundance primarily by affecting transcription rather than mRNA decay. Moreover, this relationship is not affected by optimal codon usage, although codon usage plays important roles in regulation of transcription, mRNA decay, and translation. We found that genes with high mRNA secondary structure were enriched for functions in cell maintenance, whereas genes with low mRNA secondary structure were enriched for assembly of cellular components relevant to stress response. Together, our results suggest that cells may use mRNA secondary structure, coupled with sequence features, to quickly regulate protein homeostasis and function in responses to changes in environmental conditions.

## Materials and Methods

### Yeast genomic data

mRNA sequences of *Saccharomyces cerevisiae* were retrieved from the Saccharomyces Genome Database [52].

### Experimentally determined mRNA secondary structure in yeast

Parallel analysis of RNA structure (PARS) scores for 3000 genes in yeast were taken from Kertesz et al. [33].

### Yeast mRNA levels and mRNA half-life

mRNA expression levels for 5083 genes were obtained from Pelechano et al. [43] where mRNA amount were derived from the RNA-Seq data [53], which had been normalized to real units (mRNAs/cell) using the 2000 most abundant mRNAs from the GATC-PCR data [54]. mRNA half-life data for 3888 genes were obtained from Presnyak et al. [26].

### Estimation of transcription rates and RNA polymerase II density in yeast

Nascent transcription rate data for 4666 genes obtained from an averaged Genomic Run-on dataset and RNA polymersame II density data for the same genes were taken from Pelechano et al. [43].

### Protein abundance and protein half-life in yeast

Protein abundance data for 6067 genes for *S. cerevisiae* were downloaded from PaxDb [55]. Protein half-life data for 3796 genes were obtained from Christiano et al. [56].

### Translation effciency calculation in yeast

Translation efficiency (TE) was calculated as the ratio of ribosome-protected fragments (RPF) count to mRNA abundance. TE data for 5218 genes in normal condition was retrieved from McManus et al. [57]. For our analysis of stress response, TE was calculated from the data of Gerashchenko et al. [58], which report changes in RPF and mRNA abundance for 5623 genes in response to 30 minutes of oxidative stress, compared to the normal condition.

### Prediction of RNA folding energy

The minimum free energies of mRNA sequences were calculated using the Matlab rnafold function (Matlab Bioinformatics toolbox). To measure the mean local folding of a genomic sequence, we computed and averaged the folding energy of sliding windows with a length of 30 nucleotides (nt) and a step size of 10 nt; 30 is the approximate length of the ribosomes’ footprint [59].

### Evolutionary rates

Evolutionary rates (dN, dS, and dN/dS) of genes were taken from Wall et al. [60].

### Codon usage and GC content calculation

CodonW (http://codonw.sourceforge.net/) was used to estimate Codon Adaptation Index (CAI) and GC content.

### Gene ontology (GO) enrichment analysis

GO enrichment analysis was performed with DAVID [61]. Fisher’s exact test was used to test for overrepresentation of each GO term among lowly or highly folded genes.

### Statistical analysis

Spearman’s rank correlation (or partial correlation) coefficients were used to quantify relationships between variables. Standard errors of correlation coefficients were estimated using 10,000 bootstrap data samples, and Z-tests were used to test for statistical significance of individual effects or differences between effects.

### Path analysis

Path analysis was used to investigate the contribution of PARS (or folding free energy) and CAI to Central Dogma processes. We used this path analysis to estimate the total, direct, and indirect effects of each variable on each other. For example, the indirect effect between PARS and mRNA abundance is mediated through transcription rate, CAI, and mRNA half-life. The analysis was performed using the R package sem [62].

## Results

### mRNA structured is correlated with codon usage

It has been reported that mRNA secondary structure, as measured by PARS score, is strongly associated with optimal codon usage, averaging over the gene [29]. Consistent with this report, we also found that the average PARS score over a gene is strongly positively correlated with that gene’s codon adaptation index (CAI) (Spearman rank correlation r=0.49, N=2925, p<10^−177^), a widely used index for quantifying codon usage bias [39].

To infer more specifically which codons are most strongly associated with mRNA structure, we separately analyzed all 64 codons. We found that the frequencies of most codons (49/64) are significantly correlated with gene-average PARS score (p<0.01), even after controlling for mRNA abundance and GC content (Table S1). In particular, 6 codons (ACC, GCC, GCU, GGU, GUC, UCC) are strongly positively correlated with PARS score (r>0.3, p<10^−5^), and 6 codons (AAA, AGG, AUA, CGA, GGA, GUA) are strongly negatively correlated with PARS score (r<−0.3, p<10^−5^) (Figure 1A). Codons that are positively correlated with PARS tend to be common codons with high translation efficiency [39, 40], whereas codons that are negatively correlated tend to be rare codons with low translation efficiency (Table S1). For example, both AUA and AUC code for isoleucine, and the correlation between frequency of AUA (which has low relative synonymous codon usage [39]) and PARS is r=−0.433 and between frequency of AUC (which has high relative synonymous codon usage [39]) and PARS is r=0.25. The association of mRNA secondary structure with high translation efficiency codons may act to smooth rates of translation, reducing the risk of potentially deleterious slow or fast ribosome movement [41].

**Figure 1.**
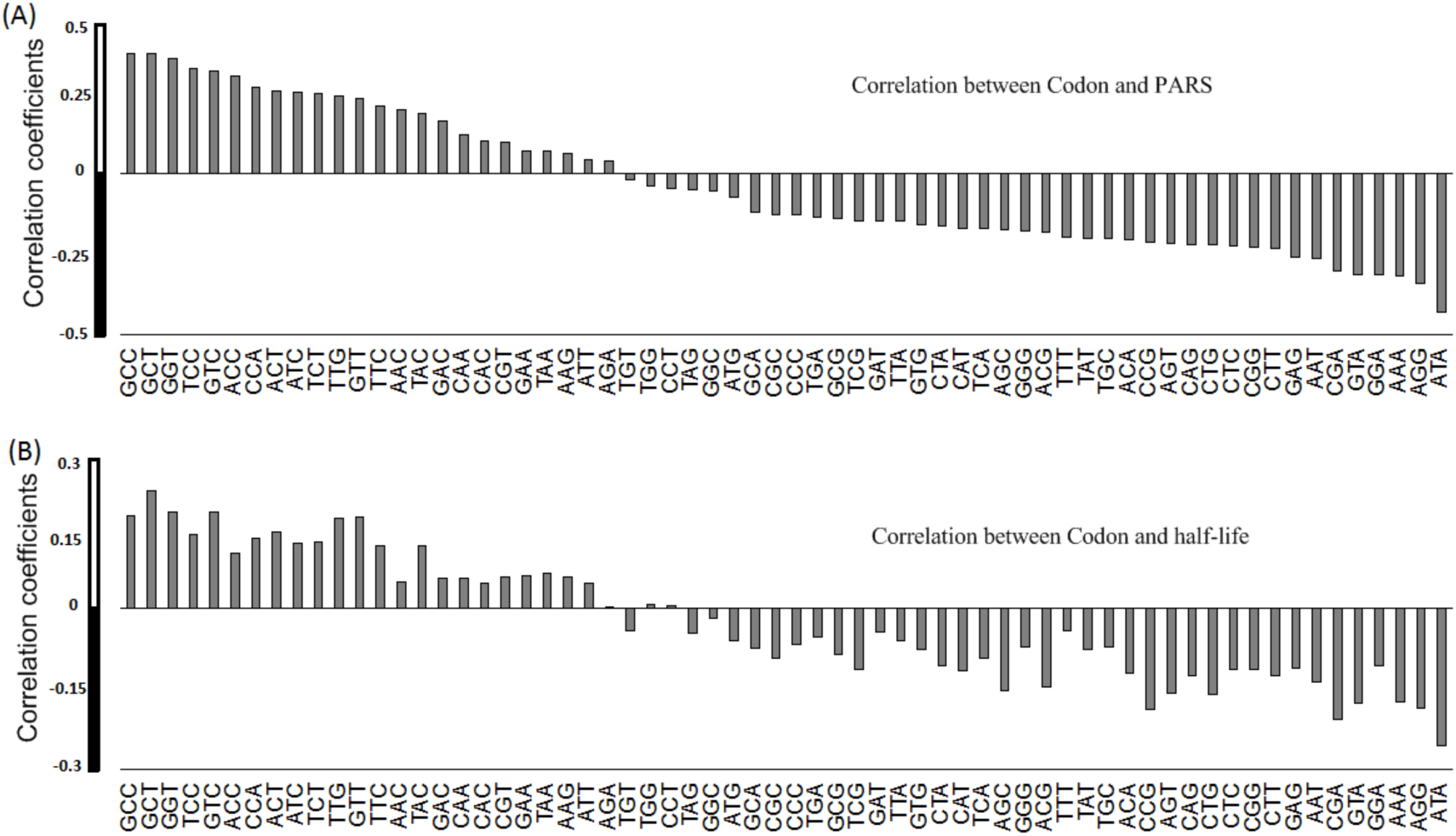
Rank correlations between frequencies within genes of all 64 codons and (A) mRNA secondary structure, measured by average PARS within each gene or (B) mRNA half-life.

In addition, the correlations between PARS score and evolutionary rates (i.e., dN, dS, and dN/dS) are negative and strong, even when we control for mRNA abundance (r=−0.29 for dS, p<0.001), suggesting that mRNAs with stronger secondary structure will evolve more slowly, potentially due to selection to preserve structure. This result is consistent with recent work of Park et al. [42], which demonstrated that mRNA folding is associated with slow evolution of highly expressed proteins.

### Contributions of mRNA secondary structure and codon bias to mRNA abundance

Previous work [30] has shown that gene-average PARS score is strongly and positively associated with mRNA abundance, an observation we corroborate (Spearman rank correlation r=0.45, N=2610, p<10^−127^; Figure S1). It is unknown, however, to what degree this correlation is driven by transcription rate and/or mRNA half-life.

To disentangle the effects of mRNA structure and other factors on gene expression, we used path analysis, a general technique for quantifying the directed acyclic dependencies among a set of variables. Path analysis begins with an assumed model, representing potential causal influences between the variables in the data. Multiple regression of the data on that model then reveals the strength of different causal links between variables. Within a given model, path analysis can thus identify the most important casual paths (those with the largest path coefficients), thus identifying the most important factors in determining the output of a complex system. Our model (Figure 2) began with the Central Dogma steps in gene expression. To those, we added potential direct influences from RNA structure (quantified as PARS) on transcription [9-12, 14], mRNA half-life [15] and translation [18, 19]. We also added potential direct influences from RNA sequence features (quantified by CAI) on transcription [23-25], mRNA half-life [24, 26], and translation [24, 27, 28]. We modeled PARS and CAI as exogenous variables, which may be correlated with each other [29] but are not causally determined by other variables in the model. The other variables, including mRNA half-life, translation, transcription, mRNA abundance, and protein abundance, are endogenous variables that depend directly or indirectly on PARS and CAI.

**Figure 2.**
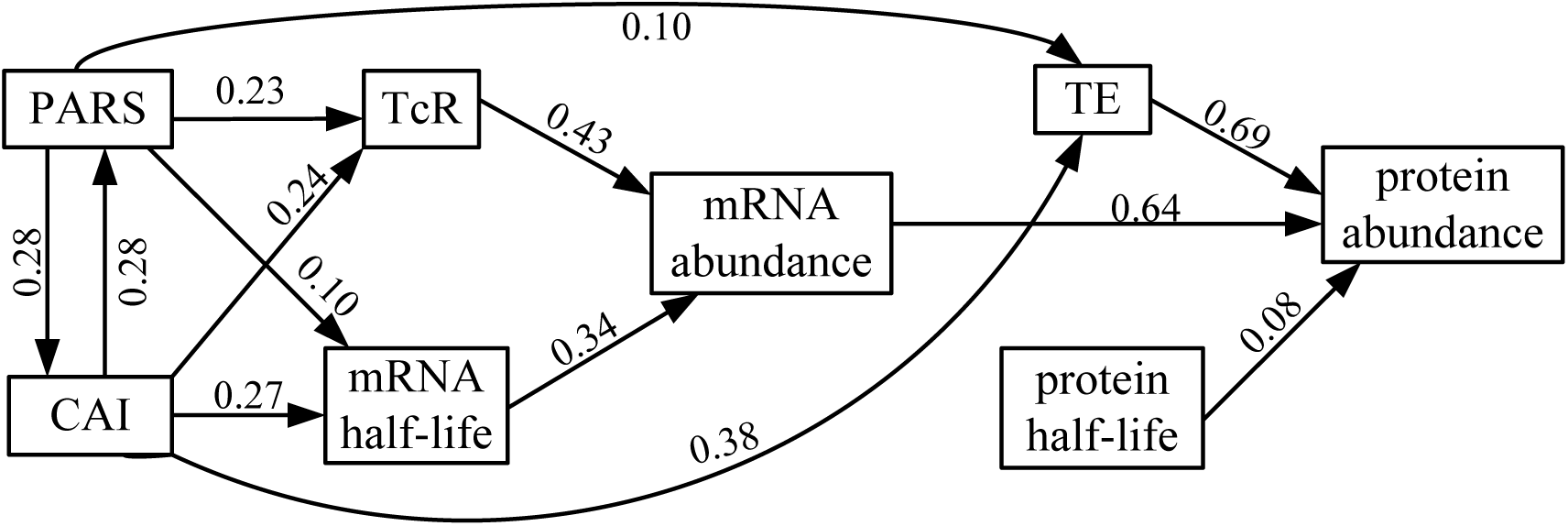
Path diagram presenting coefficients for all direct effects among mRNA secondary structure (PARS), codon adaptation index (CAI), transcription rate (TcR), mRNA half-life, mRNA abundance, translation efficiency (TE), protein half-life, and protein abundance. The potential causal relations tested between the predictor variables are based on previous investigations of relationships among secondary structure, codon bias, and transcription [9-12, 14, 23, 25], mRNA decay [15, 24, 26] and translation [18, 19, 24, 27, 28].

We applied our model to 1,344 genes for which we had data on all variables considered. We found that PARS has a larger direct effect on transcription (β=0.23) than on mRNA half-life (β=0.10; Z-test, p<0.01) (Table S2). Notably, the effect of PARS on mRNA abundance mediated by transcription (β = 0.23×0.43 = 0.099) is significantly larger than that mediated by mRNA half-life (β = 0.10×0.34 = 0.034) (Z-test, p<10^−4^). The effect of RNA structure on mRNA abundance is thus primarily driven by changes in transcription rather than mRNA half-life.

PARS is measured *in vitro*, and RNA may refold prior to analysis, which might bias our results. We thus repeated our analysis using an independent measure of RNA folding, by using a thermodynamic model to predict the minimum free energy (MFE) of each folded RNA [42]. Similar to our analysis with PARS, we found that the association between MFE and mRNA abundance mediated by transcription (β = 0.21×0.47 = 0.099) is much stronger than the association mediated by mRNA half-life (β = 0.12×0.3 = 0.036) (Z-test, p<10^−4^) (Figure S2). This provides further evidence that RNA structure affects mRNA abundance primarily through transcription and not mRNA half-life.

In contrast to PARS, CAI has similar direct effects on transcription rate (TcR) and mRNA half-life (0.24 vs. 0.27; Z-test, p=0.37). Notably, the direct effects of PARS and CAI on TcR are similar (0.23 vs. 0.24; Z-test, p=0.43) and much stronger than any indirect effects, suggesting that both RNA structure and codon content are important to transcription rate. By contrast, the effect of CAI on mRNA half-life is significantly larger than that of mRNA structure (0.24 vs. 0.10; Z-test, p<0.01). The 12 codons (ACC, GCC, GCU, GGU, GUC, UCC, AAA, AGG, AUA, CGA, GGA, GUA) most strongly correlated with PARS also have moderate correlation with mRNA stability (Figure 1B). Codon usage may thus mediate the association between PARS score and mRNA half-life. Indeed, path analysis showed that the effect of PARS on mRNA half-life as mediated by CAI (β = 0.28×0.27 = 0.076) is comparable to the direct effect of PARS on mRNA half-life (β = 0.10; Z-test, p=0.36).

Together, these analyses suggest that the correlation between mRNA structure (PARS) and mRNA abundance is driven primarily by effects of mRNA structure on transcription rate, rather than mRNA half-life. By contrast, codon usage has similar effects on transcription rate and mRNA half-life.

### Contributions of mRNA secondary structure and codon bias to translation

PARS score (r=0.51, N=2974, p<10^−193^) [30] and optimal codon usage (CAI) (r=0.50, N=5077, p<10^−178^) [28] are known to be strongly positively associated with protein abundance. Because PARS and CAI are both properties of the mRNA, they are not expected to have direct causal effects on protein half-life, so we focus on the effects of PARS and CAI on translation efficiency. Our path analysis (Figure 2) shows that the direct effect of CAI on translation efficiency is much stronger than that of PARS (0.38 vs. 0.10; Z-test, p<0.0001). The indirect effect of PARS on translation efficiency as mediated by CAI (β = 0.28×0.38 = 0.11) is similar to the direct effect (β = 0.10; Z-test, p=0.28). Codon usage thus plays a substantial role in mediating the association between PARS score and translation efficiency, and it has a much larger direct effect than PARS.

### mRNA secondary structure is associated with protein biological function

As shown above, mRNA secondary structure is significantly associated with transcriptional and translational processes. To examine the functional consequences of these associations, we asked whether there are relationships between mRNA secondary structure and biological function of the encoded proteins.

We first defined mRNAs to be highly-folded if they had an average PARS score greater than twice the average and lowly-folded if they had an average PARS score smaller than half the average. Gene Ontology (GO) enrichment analysis revealed that strongly folded transcripts were significantly (p<10^−6^) enriched for biological functions related to macromolecular (such as protein and lipid) metabolism and biosynthesis (Table 1). In contrast, weakly folded transcripts were significantly (p<10^−6^) enriched for cellular components such as mitochondrion, endoplasmic reticulum (ER) and ribosome assembly. Because PARS score is correlated with gene expression level [30], we also considered GO enrichment when genes were classified as highly- or lowly-folded while controlling for mRNA abundance (Supplemental Figure S1). Notably, we still found that the functions of lowly-folded transcripts were enriched in mitochondrion and ribosome assembly, while highly-folded transcripts were enriched in cell wall organization and metabolism-related enzyme activity (Supplemental Table S3). This suggests that highly-folded and lowly-folded transcripts contribute to different cellular processes (see Discussion).

**Table 1.** Top 30 overrepresented GO terms in sets of highly and lowly folded mRNA.

| highly folded mRNA |  |  | lowly folded mRNA |  |  |
| --- | --- | --- | --- | --- | --- |
| GO-ID | Term | p-value | GO-ID | Term | p-value |
| GO:0006417 | regulation of translation | 2.89E-33 | GO:0044429 | mitochondrial part | 2.05E-16 |
| GO:0010608 | posttranscriptional regulation of gene expression | 1.06E-32 | GO:0003735 | structural constituent of ribosome | 1.44E-14 |
| GO:0032268 | regulation of cellular protein metabolic process | 7.70E-31 | GO:0005761 | mitochondrial ribosome | 5.86E-14 |
| GO:0044445 | cytosolic part | 9.11E-31 | GO:0000313 | organellar ribosome | 5.86E-14 |
| GO:0005829 | cytosol | 1.61E-24 | GO:0033279 | ribosomal subunit | 9.42E-14 |
| GO:0022626 | cytosolic ribosome | 3.49E-20 | GO:0031090 | organelle membrane | 7.21E-13 |
| GO:0006412 | translation | 5.65E-20 | GO:0031974 | membrane-enclosed lumen | 3.97E-12 |
| GO:0005783 | endoplasmic reticulum | 7.88E-18 | GO:0030529 | ribonucleoprotein complex | 1.29E-11 |
| GO:0001950 | plasma membrane enriched fraction | 2.01E-16 | GO:0005840 | ribosome | 1.36E-11 |
| GO:0005626 | insoluble fraction | 1.55E-14 | GO:0070013 | intracellular organelle lumen | 2.82E-10 |
| GO:0005624 | membrane fraction | 1.55E-14 | GO:0043233 | organelle lumen | 2.82E-10 |
| GO:0005576 | extracellular region | 2.89E-14 | GO:0031975 | envelope | 1.26E-09 |
| GO:0022625 | cytosolic large ribosomal subunit | 1.88E-13 | GO:0031967 | organelle envelope | 1.26E-09 |
| GO:0000267 | cell fraction | 2.06E-13 | GO:0034470 | ncRNA processing | 1.81E-09 |
| GO:0009277 | fungal-type cell wall | 1.59E-12 | GO:0009451 | RNA modification | 5.20E-09 |
| GO:0008610 | lipid biosynthetic process | 8.30E-12 | GO:0005198 | structural molecule activity | 6.52E-09 |
| GO:0042802 | identical protein binding | 9.13E-12 | GO:0005759 | mitochondrial matrix | 8.40E-09 |
| GO:0005618 | cell wall | 2.02E-11 | GO:0031980 | mitochondrial lumen | 8.40E-09 |
| GO:0030312 | external encapsulating structure | 2.02E-11 | GO:0005762 | mitochondrial large ribosomal subunit | 9.42E-09 |
| GO:0043094 | cellular metabolic compound salvage | 4.88E-11 | GO:0000315 | organellar large ribosomal subunit | 9.42E-09 |
| GO:0006696 | ergosterol biosynthetic process | 1.86E-10 | GO:0008033 | tRNA processing | 1.07E-08 |
| GO:0016129 | phytosteroid<br>biosynthetic<br>process | 1.86E-10 | GO:0042254 | ribosome biogenesis | 1.56E-08 |
| GO:0000502 | proteasome<br>complex | 3.04E-10 | GO:0012505 | endomembrane<br>system | 2.26E-08 |
| GO:0046394 | carboxylic acid<br>biosynthetic<br>process | 3.38E-10 | GO:0005740 | mitochondrial<br>envelope | 2.81E-08 |
| GO:0016053 | organic acid<br>biosynthetic<br>process | 3.38E-10 | GO:0044432 | endoplasmic<br>reticulum part | 3.01E-08 |
| GO:0055114 | oxidation reduction | 4.37E-10 | GO:0031966 | mitochondrial<br>membrane | 5.33E-08 |
| GO:0044271 | nitrogen compound<br>biosynthetic<br>process | 4.44E-10 | GO:0032543 | mitochondrial<br>translation | 1.42E-07 |
| GO:0031597 | cytosolic<br>proteasome<br>complex | 1.03E-09 | GO:0015935 | small ribosomal<br>subunit | 3.05E-07 |
| GO:0034515 | proteasome storage<br>granule | 1.03E-09 | GO:0019866 | organelle inner<br>membrane | 3.14E-07 |
| GO:0016126 | sterol biosynthetic<br>process | 1.19E-09 | GO:0005743 | mitochondrial inner<br>membrane | 3.68E-07 |

## Discussion

RNA’s ability to adopt complex secondary and tertiary folds has critical roles in processes ranging from ligand sensing to the regulation of translation, polyadenylation, and splicing [21, 22]. Here we investigated the relationships between mRNA secondary structure and codon content on multiple steps of gene expression (transcription, mRNA decay, translation) on a genome-wide scale.

We found that mRNA secondary structure is strongly positively correlated with transcription rate, and path analysis suggests that this is the dominant cause of the correlation between mRNA secondary structure and mRNA abundance. How does mRNA secondary structure affect transcription rate? Previous research has shown that the net transcription rate, whether limited by initiation or elongation, is strongly correlated with the density of RNA Polymerase II on the gene [43, 44]. We speculate that co-transcriptional folding of mRNA enhances transcription by reducing interaction between the nascent mRNA and adjacent polymerases, thereby enabling increased polymerase density. This model predicts that, for a given transcription rate, longer genes will need to fold more tightly, because their mRNAs are bulkier. Indeed, we find that gene length is significantly positively correlated with PARS score (r=0.23, N=3001, p<10^−35^, Figure 3). Meanwhile, longer genes also probably have more regulating transcription factors that will also affect gene expression [45].

**Figure 3.**
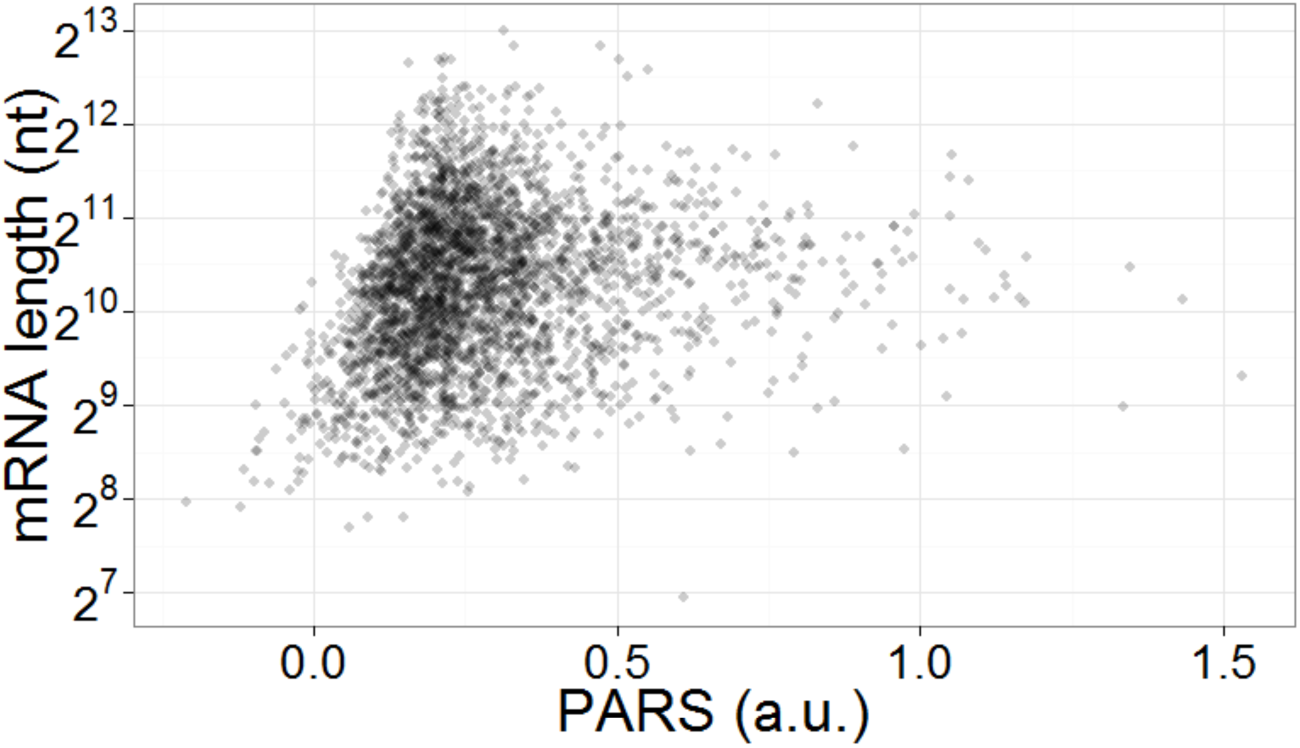
Strong correlation between mRNA length and mRNA secondary structure (PARS score).

We found that codon bias has much stronger direct effects on translation than mRNA structure, but they may still play complementary roles. Prior work suggests that local association within genes between mRNA secondary structure and high translation efficiency codons may act to smooth rates of translation [41]. We also found that strongly folded mRNAs have lower synonymous evolutionary rates dS, suggesting that selection is acting to conserve mRNA secondary structure [42, 46], thereby conserving its effects on gene expression. Together, these results suggest that a global coupled transcriptional and translational process controlling mRNA and protein abundance has strong association with mRNA secondary structure and other sequence information.

Growing evidence shows the importance of mRNA expression level in regulating transient responses to stress and stimulus such as temperature, oxidation, starvation, and metal ions [47, 48]. These stresses and stimuli can not only affect RNA structure and thermostability [47, 48], but also induce ribosomes, the endoplasmic reticulum, and mitochondria to undergo extensive structural and/or spatial reorganization [49, 50]. So are there any relationships between mRNA structure and biological functions of the encoded proteins? Our GO enrichment analysis revealed that lowly-folded mRNAs tend to function in the mitochondria and endoplasmic reticulum and in ribosome assembly. Recent *in vivo* genome-wide profiling of RNA secondary structure in *Arabidopsis thaliana* seedlings revealed that highly-folded mRNAs are related to cell function maintenance and lowly-folded mRNAs are related to stress and stimulus response [12]. Thus, lowly-folded mRNAs in yeast annotated for mitochondrion, ER, and ribosome function may also be relevant to stress response. In fact, ER ribosomes are the primary sites for synthesis and folding of proteins, which is sensitive to extracellular stresses and stimuli. Moreover, the ER and mitochondria interact both physiologically and functionally. Both are dynamic organelles capable of modifying their structure and function to maintain cellular homeostasis and determine cell fate in response to changing environmental conditions [51]. Consistent with this model, in yeast RNA secondary structure (PARS) is negatively correlated (r=−0.402, p<10^−16^) with fold change in translation efficiency in stress vs. normal conditions (Figure 4). Lowly-folded transcripts tend to have increased translation efficiency during stress condition, whereas highly-folded transcripts have reduced translation efficiency.

**Figure 4.**
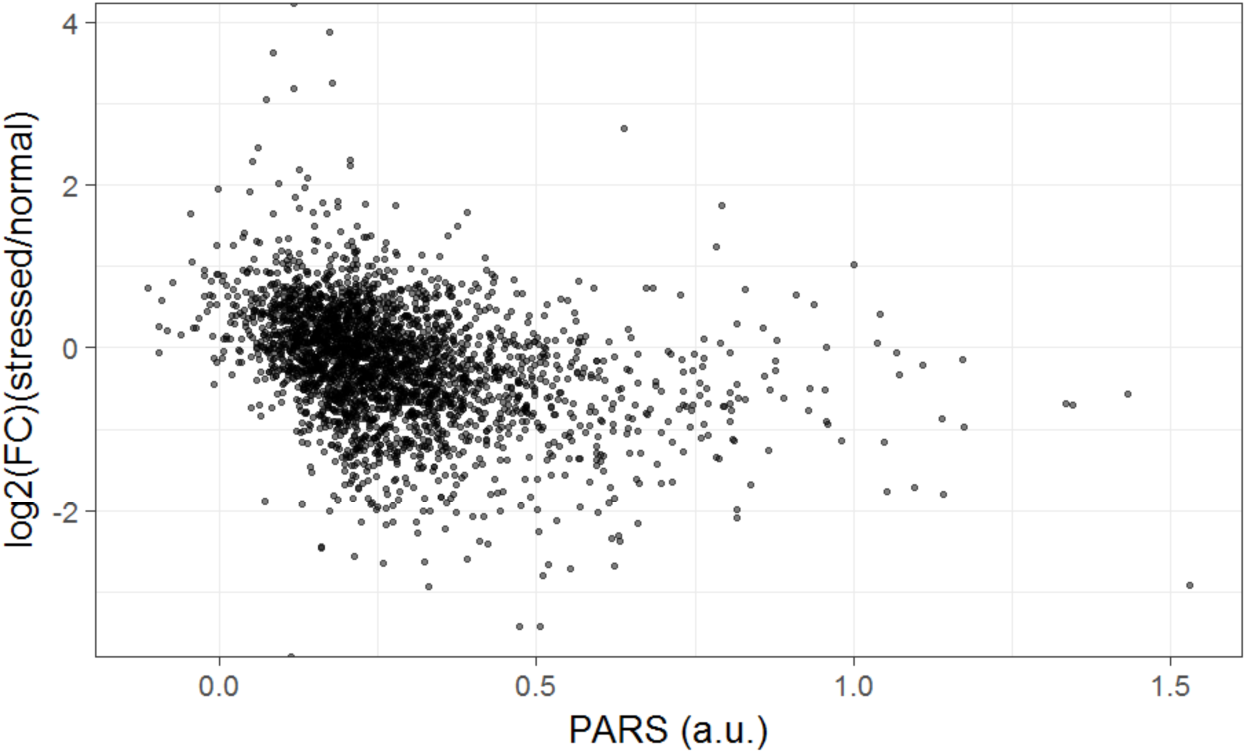
Strong correlation between fold change of translation efficiency in stress condition vs. normal condition and mRNA secondary structure (PARS score).

Combining our results, we found that lowly-folded mRNAs are related to mitochondrion, ER, and ribosome assembly and have short half-life and low transcription rates. After removal of stress, slow synthesis and rapid removal of stress-responsive mRNAs may thus facilitate rapid return of the cell to a normal state.

## Additional material

Additional file 1: Supplementary Figures S1 and S2, Tables S1, S2, and S3.

## Abbreviations

PARS: parallel analysis of RNA structure
RNAPII: RNA polymerase II
CAI: codon adaptation index
GO: Gene Ontology
ER: endoplasmic reticulum.

## Acknowledgements

This work was supported by DARPA contract WF911NF-14-1-0395 to R.G., and the NSF of China and Anhui Province (Grant No. 31500676 and 1508085SQC202, to X.W.). We thank Brian Mannakee and Travis Struck for helpful discussions.

## Authors’ contributions

XW and RG designed the analyses and wrote the manuscript. XW and PL conducted the data analysis. All authors have read and approved the manuscript for publication.

